# Ion transport mechanisms for smoke inhalation injured airway epithelial barrier

**DOI:** 10.1101/2020.03.25.007807

**Authors:** Jianjun Chang, Zaixing Chen, Runzhen Zhao, Hong-Guang Nie, Hong-Long Ji

**Affiliations:** Department of Cellular and Molecular Biology, University of Texas Health Science Center at Tyler, 11937 US Hwy 271, Tyler Texas 75708, USA; Institute of Health Sciences, China Medical University, Shenyang 110122, Liaoning, China; Department of Stem Cells and Regenerative Medicine, College of Basic Medical Science, China Medical University, Shenyang 110122, Liaoning, China; Texas Lung Injury Institute, University of Texas Health Science Center at Tyler, Tyler Texas 75708, USA

**Keywords:** thermal stress, acrolein, tracheal epithelial monolayers, ion transport, tight junctions

## Abstract

Smoke inhalation injury is the leading cause of death in firefighters and victims. Inhaled hot air and toxic smoke are the predominant hazards to the respiratory epithelium. We aimed to analyze the effects of thermal stress and smoke aldehyde on the permeability of the airway epithelial barrier. Transepithelial resistance (R_TE_) and short-circuit current (I_SC_) of mouse tracheal epithelial monolayers were digitized by an Ussing chamber setup. Zonula occludens-1 tight junctions were visualized under confocal microscopy. A cell viability test and fluorescein isothiocyanate-dextran assay were performed. Thermal stress (40°C) decreased R_TE_ in a two-phase manner. Meanwhile, thermal stress increased I_SC_ followed by its decline. Na^+^ depletion, amiloride (an inhibitor for epithelial Na^+^ channels [ENaCs]), ouabain (a blocker for Na^+^/K^+^-ATPase) and CFTRinh-172 (a blocker of cystic fibrosis transmembrane regulator [CFTR]) altered the responses of R_TE_ and I_SC_ to thermal stress. Steady-state 40°C increased activity of ENaCs, Na^+^/K^+^-ATPase, and CFTR. Acrolein, one of the main oxidative unsaturated aldehydes in fire smoke, eliminated R_TE_ and I_SC_. Na^+^ depletion, amiloride, ouabain, and CFTRinh-172 suppressed acrolein-sensitive I_SC_, but showed activating effects on acrolein-sensitive R_TE_. Thermal stress or acrolein disrupted zonula occludens-1 tight junctions, increased fluorescein isothiocyanate-dextran permeability but did not cause cell death or detachment. The synergistic effects of thermal stress and acrolein exacerbated the damage to monolayers. In conclusion, the paracellular pathway mediated by the tight junctions and the transcellular pathway mediated by active and passive ion transport pathways contribute to impairment of the airway epithelial barrier caused by thermal stress and acrolein.

**Graphical Headlights:**

- Thermal stress and acrolein are two essential determinants for smoke-inhalation injury, impairing airway epithelial barrier.
- Transcellular ion transport pathways via the ENaC, CFTR, and Na/K-ATPase are interrupted by both thermal stress and acrolein, one of the most potent smoke toxins.
- Heat and acrolein damage the integrity of the airway epithelium through suppressing and relocating the tight junctions.

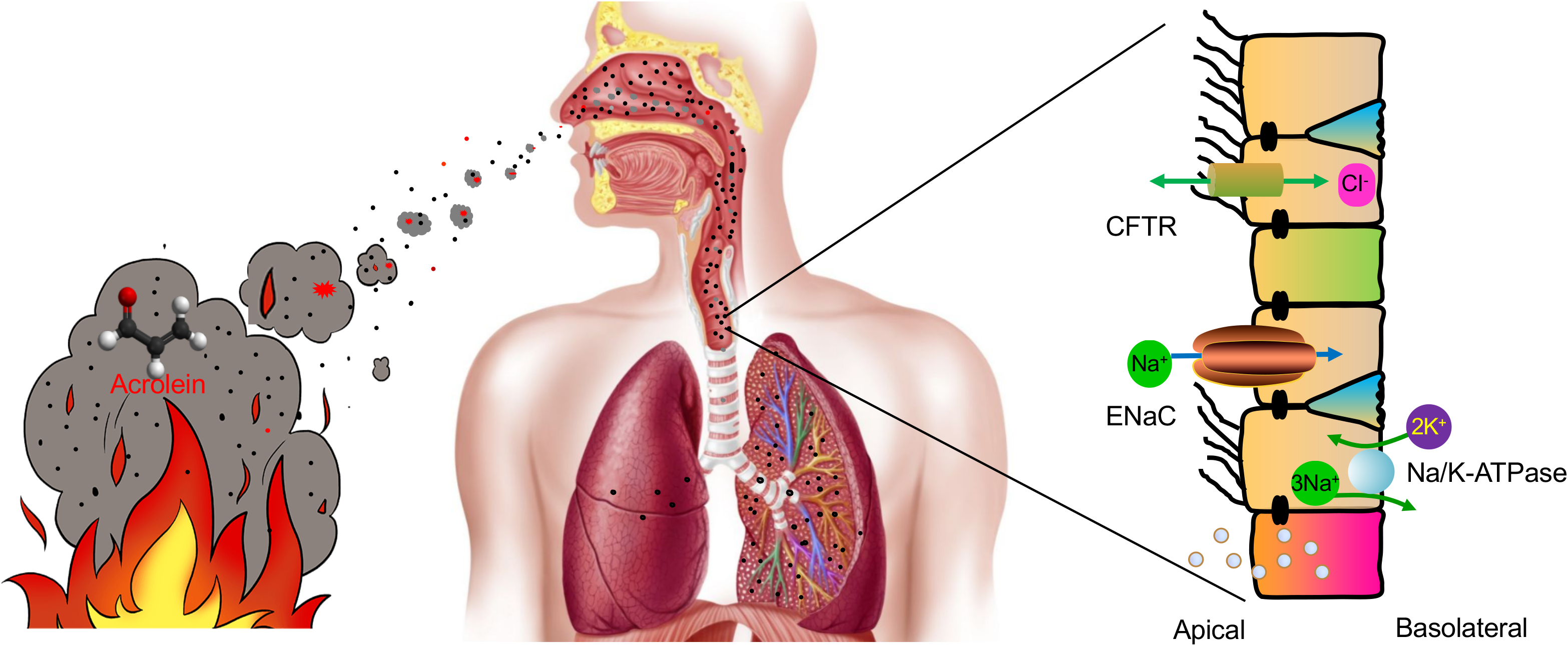

## Introduction

Fire is a common disaster in urban and rural populations. Smoke is a complicated mixture of hot air, toxic chemicals, and suspended particles (Fitzgerald and Flood 2006). Inhalation of harmful smoke causes direct impairment of respiratory function and is the leading cause of fire-related death (Haponik 1993). Inhaled hot air and toxic chemicals in smoke from fire impair the respiratory tract and lungs.

The airway epithelium is the most important defense barrier in the respiratory tract (Wang et al. 2008). Approximately 80% of the cells are ciliated in the normal respiratory tract. These cells form a tight airway epithelial layer, which can effectively prevent various external attacks (Hou et al. 2019). Destruction of the integrity of the airway epithelial barrier can be an important indicator of occurrence and the development of many respiratory diseases (Wang et al. 2007). Under pathological conditions, the integrity of the airway epithelial barrier is disrupted and its permeability is increased. Permeability of a tight airway epithelium is determined by paracellular and transcellular pathways. Paracellular pathway is predominantly regulated by tight junctions and lateral intercellular space. Transcellular permeability is predominantly controlled by active and passive ion transport pathways. Apical epithelial Na^+^ channels (ENaCs) and cystic fibrosis transmembrane regulator (CFTR), as well as basolateral Na^+^/K^+^-ATPase, are mainly responsible for Na^+^/Cl^−^ transport in the respiratory epithelium (Chang et al. 2018; Wang et al. 2007). In respiratory epithelium, passive ion transport mediated by ENaCs is an important determinant of transcellular permeability, and active Na^+^/K^+^-ATPase provides the driving force for passive ion transport systems.

A previous study showed the mats of ciliated cells in rat tracheal epithelium post-thermal injury (Dubick et al. 2002). Thermal stress regulates ENaCs in alveolar epithelial cells (Howard et al. 2013) and CFTR is implicated in thermal stress-induced signaling events in bronchial epithelial cells (Dong et al. 2015). Additionally, Na^+^/K^+^-ATPase is heat-sensitive in rabbit esophageal epithelium (Tobey et al. 1999). However, whether thermal stress regulates permeability of the airway epithelial barrier through the paracellular pathway (mediated by tight junctions) and the transcellular pathway (mediated by ion channels and transporters) in primary airway epithelial cells remains unclear.

Smoke aldehydes are released from the burning of wood and paper products, as well as cotton fabrics. Formaldehyde and acrolein are the two main gaseous α, β unsaturated aldehydes in fire and cigarette smoke (Reinhardt and Ottmar 2004; Anthony et al. 2007; Alwis et al. 2015). Inhalation of formaldehyde and acrolein results in gene induction, inflammation, cell apoptosis, and necrosis in the respiratory epithelium (Bein and Leikauf 2011; Meacher and Menzel 1999). Our previous studies showed that formaldehyde and crotonaldehyde inhibited the activity and expression of ENaCs in alveolar epithelial cells (Cui et al. 2016; Li et al. 2017). Acrolein led to irritation of the airway and impaired fluid homeostasis (Borchers et al. 1998; Romet-Haddad et al. 1992; Roux et al. 2002), as well as downregulated CFTR activity in respiratory epithelial cells (Alexander et al. 2012). Acrolein is used to prepare animal models for acute lung injury and lung edema (Hales et al. 1988). However, the pathological role of acrolein in the presence of thermal stress in the smoke-injured airway epithelial barrier has not been studied.

Therefore, this study aimed to investigate the effects of thermal stress or/and acrolein on permeability of the airway epithelial barrier. We recorded the transepithelial resistance (R_TE_) and short-circuit current (I_SC_) in primary mouse tracheal epithelial (MTE) monolayers in an Ussing chamber setup. To further determine the underlying transcellular mechanisms, inhibitors for ion channels and transporters (i.e., amiloride, ouabain, and CFTRinh-172) were applied to MTE monolayers. To examine the role of the paracellular pathway, the tight junctions were visualized under confocal microscopy, and the fluorescein isothiocyanate (FITC)-dextran assay was performed.

## Materials and methods

### Animals

Forty healthy wild-type C57BL/6 mice (males, 20; females, 20), 8-12 weeks old (mean: 10.86 ± 0.28 weeks) and weighed 18-25 g (mean: 20.03 ± 0.30 g), were purchased from the Jackson Laboratory and the Laboratory Animal Center of China Medical University. Mice were housed in a pathogen-free facility, the husbandry condition of which included a suitable light/dark cycle, temperature, drinking water, and food. All experiments were performed according to the guidelines and regulations of the Animal Care and Use Ethics Committee, and all protocols were approved by the University of Texas Health Science Center at Tyler and China Medical University. The reference numbers of ethics approval are IACUC 611 and SCXK (Liao) 2018-0001.

### Isolation and culture of MTE cells

Pooled MTE cells were isolated from C57BL/6 mice and cultured according to our previous study (Chen et al. 2014). Briefly, tracheas of anesthetized mice were removed, and cleaned tracheas were incubated in Dulbecco’s modified eagle medium (cat# 30-2002, American Type Culture Collection, USA) containing 0.1% protease XIV (cat# P5147, Sigma, USA), 0.01% DNase (cat #DN25, Sigma-Aldrich, St. Louis, MO), and 1% fetal bovine serum (cat# 26140-087, Gibco, USA) at 4°C for 24 h. MTE cells were seeded onto 6.5-mm diameter, collagen IV (cat# 3410-010-01, Trevigen, Gaithersburg, MD)-coated transwell inserts (cat# 3413, Corning-Costar, Lowell, MA) at a density of 3.0 × 10^5^/cm^2^. These cells were grown in a 1:1 mixture of Ham’s F-12 medium (cat# 11765-054, Invitrogen, Camarillo, CA) and 3T3 fibroblast preconditioned Dulbecco’s modified eagle medium supplemented with insulin (10 μg/ml, cat# I1882, Sigma-Aldrich, St. Louis, MO), hydrocortisone (1 μM, cat# H0396, Sigma-Aldrich, St. Louis, MO), endothelial cell growth supplement (3.75 μg/ml, cat# E0760, Sigma-Aldrich, St. Louis, MO), epidermal growth factor (25 ng/ml, cat# E4127, Sigma-Aldrich, St. Louis, MO), triiodothyronine (30 nM, cat# T6397, Sigma-Aldrich, St. Louis, MO), iron-saturated transferrin (5 μg/ml, cat# T1283, Sigma-Aldrich, St. Louis, MO), cholera toxin (10 ng/ml, cat# C8052, Sigma-Aldrich, St. Louis, MO), and dexamethasone (250 nM, cat# D2915, Sigma-Aldrich, St. Louis, MO). MTE cells from one mouse were seeded onto three to four transwell inserts and cultured for up to 12 days. Polarized monolayers with a reading of R_TE_ >1000 Ω by an epithelial voltohmmeter (WPI, Sarasota, FL) were used. Monolayers from the same batch of mice were randomly allocated to control and treated groups.

### Preparation of human bronchial epithelial monolayers

Human bronchial epithelial (HBE) cells were purchased from American Type Culture Collection. Cells were seeded in plastic T-75 flasks and grown in keratinocyte culture medium (cat# 2101, Sciencell, China) supplemented with keratinocyte growth factor (1%). The culture medium was changed every 48 h until 90% confluent. HBE cells were seeded (10^6^ cells/cm^2^) onto collagen IV-coated transwell inserts and grown in keratinocyte culture medium supplemented with keratinocyte growth factor (1%), insulin-transferrin-selenium solution, (1.5%, cat# 41400045, Gibco, USA) and dexamethasone (1 μM, cat# D2915, Sigma-Aldrich, St. Louis, MO). Polarized monolayers at 12-day post-seeding were used. At all stages of culture, cells were maintained at 37°C in 5% CO2 in an air incubator.

### Preparation and administration of ion transport regulators

Amiloride (an inhibitor for ENaCs, cat# A4562, Sigma-Aldrich, St. Louis, MO) was reconstituted in an apical bath solution containing 50% dimethyl sulfoxide (DMSO). CFTRinh-172 (an inhibitor for CFTR, cat# C2992, Sigma-Aldrich, St. Louis, MO) was reconstituted in DMSO. Ouabain (an inhibitor for Na^+^/K^+^-ATPase, cat# O3125, Sigma-Aldrich, St. Louis, MO) was reconstituted in a basolateral bath solution. Reconstituted amiloride (100 μM) or CFTRinh-172 (20 μM) was added to the apical bath solution of MTE monolayers. Reconstituted ouabain (1 mM) was added to the basolateral bath solution. Vehicle solution of an equal volume was added to the corresponding bath solution as the control.

### Concurrent measurements of I_SC_ and R_TE_

Measurement of I_SC_ and R_TE_ of MTE or HBE monolayers was performed as we described previously (Han et al. 2010; Nie et al. 2009; Li et al. 2017). In brief, monolayers were mounted into Ussing chambers (Physiologic Instruments, San Diego, CA) and *bathed in* saline solution containing (in mM) 120 NaCl, 25 NaHCO_3_, 3.3 KH_2_PO_4_, 0.83 K_2_HPO_4_, 1.2 CaCl_2_, 1.2 MgCl_2_, 10 HEPES, 10 D-mannitol (apical), and 10 D-glucose (basolateral). The saline bath solution was bubbled continuously with a gas mixture of 95% O_2_ and 5% CO_2_. Transepithelial potential was short-circuited to 0 mV, and R_TE_ and I_SC_ levels were measured with an epithelial voltage clamp (VCCMC8, Physiologic Instruments, San Diego, CA). A 10-mV pulse of 1 s duration was imposed every 10 s to monitor R_TE_ levels. When R_TE_ and I_SC_ levels were stable for at least 10 min, monolayers were treated as follows: 1) amiloride (100 μM) was added to the apical bath solution; 2) CFTRinh-172 (20 μM) was added to the apical bath solution; 3) ouabain (1 mM) was applied to the basolateral bath solution; and 4) saline bath solution of both apical and basolateral chambers was replaced with Na^+^-free bath solution or Cl^−^-free bath solution. The same volume of vehicle solution was added to corresponding chambers, which was used as the control of amiloride, CFTRinh-172, or ouabain. The composition of the Na^+^-free bath solution contained (in mM) 120 N-methyl-D-glucamine-Cl, 25 KHCO3, 3.3 KH_2_PO_4_, 0.83 K_2_HPO_4_, 1.2 CaCl_2_, 1.2 MgCl_2_, 10 HEPES, 10 D-mannitol (apical), and 10 D-glucose (basolateral). The Cl^−^-free bath solution contained (in mM) 120 sodium gluconate, 25 NaHCO_3_, 3.3 KH_2_PO_4_, 0.83 K_2_HPO_4_, 1.2 Ca(NO3)2, 1.2 MgSO4, 10 HEPES, 10 D-mannitol (apical), and 10 D-glucose (basolateral). Before replacement of Na^+^-free or Cl^−^-free bath solution, recording of R_TE_ and I_SC_ in the Ussing chamber system was stopped, and then saline bath solution was aspirated from the apical and basolateral chambers. Warm saline (as the control), Na^+^-free bath solution, or Cl^−^-free bath solution was then added to the chambers. Finally, recording in the Ussing chamber system was restored.

### Application of thermal stress or acrolein to monolayers

All experiments of thermal stress or acrolein application were simultaneously performed with recording in the Ussing chamber system. In the experiments for studying the effects of thermal stress on R_TE_ and I_SC_, and the underlying mechanisms, monolayers were mounted into the chambers in which the temperature was 37°C. The temperature of bath solution of the apical and basolateral chambers was gradually elevated to the designed degree (40°C) from 37°C. A bath circulator (Thermo Fisher Scientific, CA) was used to regulate the temperature of the chambers. The bath circulator was set to the desired temperature. Water was pumped from the bath of the circulator and continuously recirculated around the chambers, thereby transferring heat to the bath solution in the chambers. Additionally, thermometers in the apical and basolateral bath solution of the chambers were used to monitor and confirm the temperature setting. In the experiments for investigating the effects of thermal stress on ion transport system, the bath solution in the chambers was first set to the desired temperature (40°C), and then monolayers were mounted into chambers to achieve instant exposure of thermal stress. In the experiments of acrolein exposure, a range of working concentrations from 5 to 500 μM was accumulatively pipetted into the apical bath solution. Acrolein vapor was minimized without gas bubbling of the bath solution, and the bath solution was moderately blown up to 10 times to ensure even distribution of acrolein. To study the synergistic effects of acrolein and thermal stress, monolayers were mounted into chambers in which the temperature was already 40°C or 37°C (as the control). When R_TE_ and I_SC_ traces were stable, a series of concentrations of acrolein were added to the apical bath solution.

### Immunofluorescent staining and imaging of MTE monolayers

MTE monolayers that were *bathed in* saline solution in Ussing chambers were exposed to 40°C (120 min) or acrolein (500 μM). For synergistic treatment of thermal stress and acrolein, MTE monolayers were *bathed in* saline solution (40°C or 37°C), and then 500 μM acrolein was added to the apical bath solution. After this treatment, monolayers were prepared for staining. Monolayers were fixed in phosphate-buffered saline containing 4% paraformaldehyde for 15 min and washed with phosphate-buffered saline. Monolayers were permeabilized with 0.5% Triton X-100 and blocked with a blocking buffer containing 10% goat serum (cat# G9023, Sigma-Aldrich, St. Louis, MO) and 1% bovine serum albumin (cat# A3803, Sigma-Aldrich, St. Louis, MO) for 1 h. Monolayers were incubated with rabbit anti-mouse zonula occludens-1 (ZO-1) antibody (1:50; cat# 61-7300, Invitrogen, Camarillo, CA) or rabbit anti-mouse occludin antibody (1:200; cat# 40-4700, Invitrogen, Camarillo, CA) in blocking buffer overnight at 4°C. Cells were incubated with Alexa Fluor 488-labeled goat anti-rabbit IgG (1:1000, cat# 111-545-045 Jackson Immuno Research Laboratories, Carlsbad, CA) and Hoechst (1:1000, cat# 561908, BD Biosciences, San Jose, CA) and then mounted with Vectashield mounting media (cat# H1000, Vector Laboratories, Burlingame, CA). Images were obtained with a Zeiss LSM 510 confocal microscope (Carl Zeiss AG, Germany). A series of optical sections were collected at 1-μm intervals in the Z-axis. Images and quantification of fluorescence intensity were analyzed with ImageJ (NIH).

### Cell counting kit-8 assay

Cell viability was quantified by the cell counting kit-8 assay (cat# C0037, Beyotime, Jiangsu, China). Culture medium containing 10% cell counting kit-8 solution was added apically to MTE monolayers at post-treatment of 37°C (control), 40°C (120 min), acrolein (500 μM), or 40°C combined with acrolein. After 2 h of incubation at 37°C, culture medium in transwell inserts was collected. Absorbance was measured at a wavelength of 450 nm.

### FITC-dextran assay

To measure dextran permeability, FITC-dextran (4 kDa, 1 mg/ml, cat# FD4, Sigma-Aldrich, St. Louis, MO) was added to the transwell inserts. Aliquots were withdrawn from the lower chambers after 4 h and assayed for fluorescence at 530 nm.

### Statistical analysis

Data are expressed as the mean ± SEM. Normality and homoscedasticity tests were performed by the Shapiro-Wilk and Levene tests. The real power of the sample size with alpha = 0.05 is shown in the figure legends. The student’s two-tailed t-test was used for comparing two groups with parametric data. For comparison of multiple groups, we performed one-way analysis of variance followed by Bonferroni’s test for all groups in the experiments. For non-parametric data, the Mann-Whitney U test was used. Differences were considered significant when the *P* value was < 0.05. The τ_1/2_ value was computed by fitting R_TE_ raw data with the ExpDec1 function (*y* = *y*_0_ + *Ae^-x/t^*), where *y* is R_TE_, *x* is time, *y*_0_ is offset; and *A* is amplitude. The dose-response curve for acrolein was generated by fitting the raw data with the Hill equation (*y = START* + (*END-START*)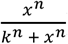), where *y* is the normalized R_TE_/I_SC_ value, *x* is the logarithmic concentration of acrolein, *START* and *END* are initial and ending values of R_TE_, respectively, *k* is the *Ki* value, and *n* is the total number of *x*. In thermal stress-treated R_TE_/I_SC_ traces, ΔR_TE_ was defined as the difference between the initial R_TE_ and end of R_TE_ within one phase (phase 1 [P1] or phase 2 [P2]) or the entire procedure, and ΔI_SC_ was defined as the difference between the basal I_SC_ and the peak I_SC_ at P1 or the difference between the peak I_SC_ and end of I_SC_ at P2. In acrolein-treated R_TE_/I_SC_ traces, ΔR_TE_/ΔI_SC_ was defined as the difference between basal R_TE_/I_SC_ and acrolein-resistant R_TE_/I_SC_. ENaCs, CFTR, or Na^+^/K^+^-ATPase activity was determined as the difference between the basal I_SC_ and corresponding specific inhibitor-resistant I_SC_. Statistical analysis was performed with Origin Pro 8.0 (OriginLab Corp. MA).

## Results

### Thermal stress alters R_TE_ and I_SC_ in MTE and HBE monolayers

The airway epithelial barrier is the primary line of defense against external attacks. Under pathogenic conditions, the integrity of this barrier is damaged and permeability is increased. We postulated that thermal stress would regulate permeability of the airway epithelium. In our study, basal R_TE_ and I_SC_ in polarized MTE monolayers were recorded at 37°C. To investigate the effect of thermal stress, the temperature of the bath solution was elevated from 37°C. R_TE_ and I_SC_ were eliminated in minutes when the temperature of the bath solution was elevated to 44°C or 42°C (Fig. S1). Therefore, 40°C was selected for the temperature of thermal stress. R_TE_ was decreased when the temperature of the bath solution was raised to 40°C from 37°C. Moreover, the entire process was divided into two stages of P1 and P2 (Fig. 1a). At each phase, R_TE_ was significantly decreased in a time-dependent manner (Fig. 1c). We used τ_1/2_ to reflect the declining rate of R_TE_ within one phase, which was computed by fitting R_TE_ raw data with the ExpDec1 function. The τ_1/2_ at P2 was significantly higher than that at P1 (Fig. 1e, *P* < 0.001). We defined P1 and P2 of I_SC_ using the R_TE_ time course. In contrast, I_SC_, which was recorded in parallel with R_TE_, was increased at P1 and decreased at P2 (Fig. 1b and 1d, all *P* < 0.01). To overcome the difference between species, we recorded R_TE_ and I_SC_ in HBE monolayers during thermal stress. We found that R_TE_ of HBE monolayers was decreased by thermal stress, which was similar to the result from MTE monolayers (Fig. 1f and 1h, *P* = 0.077). However, I_SC_ of HBE monolayers was increased to a high and stable level by thermal stress (Fig. 1g and 1i, \**P* < 0.05). Our data suggest that R_TE_ and I_SC_ of primary MTE monolayers respond to thermal stress in a phase-dependent, diverse manner.

### Na^+^ transport mediates thermal stress-induced bioelectric changes

In respiratory epithelium, Na^+^ transport is mainly regulated by ENaCs and Na^+^/K^+^-ATPase (Chang et al. 2018). Passive ion transport across tight epithelial monolayers includes ion channels (i.e., ENaCs) and transporters, which are an important determinant of transcellular permeability. Active Na^+^/K^+^-ATPase provides the driving force for the passive ion transport systems. To examine the role of epithelial Na^+^ transport in thermal stress-altered R_TE_ and I_SC_, Na^+^-free bath solution, amiloride, or ouabain was applied to MTE monolayers *bathed in* Ussing chambers at 37°C followed by elevation to 40°C. At 37°C, Na^+^-free bath solution and amiloride, but not ouabain, caused an increment in R_TE_ (Fig. 2a, black line). A drop in I_SC_ was seen at this time (Fig. 2b, black line). During thermal stress, R_TE_ was decreased in a two-phase manner in the presence of amiloride, which was similar to the result from the saline group. In contrast, a continuous decrease in R_TE_ was observed in the presence of Na^+^-free bath solution or ouabain, which could not be divided into two significant phases (Fig. 2a, red line). I_SC_ at P1 and P2 was observed in the presence of Na^+^-free bath solution or amiloride (Fig. 2b, red line). We computed thermal stress-sensitive R_TE_ at P1 and P2 (ΔR_TE_, in Fig. 2c), and found that amiloride markedly increased thermal stress-sensitive R_TE_ at P1 (*P* < 0.01), but there were no significant effects at P2 (*P* = 0.56). Thermal stress caused the continuous reduction of R_TE_ in the presence of the Na^+^-free bath solution and ouabain, so we calculated thermal stress-sensitive R_TE_ in the entire procedure (ΔR_TE_ in Fig. 2d), and found that Na^+^-free bath solution markedly increased thermal stress-sensitive R_TE_ and ouabain significantly decreased thermal stress-sensitive R_TE_ in the entire of procedure (all *P* < 0.05). We calculated thermal stress-sensitive I_SC_ at P1 and P2 (ΔI_SC_, in Fig. 2e and 2f), and found that thermal stress-sensitive I_SC_ at P1 was significantly decreased in the presence of Na^+^-free bath solution, amiloride, or ouabain (Fig. 2e, all *P* < 0.05). Additionally, Na^+^-free bath solution and amiloride markedly decreased thermal stress-sensitive I_SC_ at P2 (Fig. 2f, *P* < 0.001). Because we could not observe the obvious I_SC_ P2 in the presence of ouabain, we did not calculate thermal stress-sensitive I_SC_ at P2 in the presence of ouabain. These data indicate that changes in R_TE_ and I_SC_ caused by thermal stress may be predominately determined by transcellular Na^+^ transport powered by ENaCs and Na^+^/K^+^-ATPase. Interestingly, blockade of Na^+^/K^+^-ATPase showed lower thermal stress-sensitive I_SC_ compared with Na^+^-free bath solution. The potential mechanisms of this finding may be the temperature-dependent Na^+^/K^+^-ATPase and others, including ENaCs, CFTR, and K^+^ channels (KCs). We attempted to further confirm the effects of thermal stress on permeabilized half monolayers. However, neither R_TE_ nor I_SC_ levels were stable enough to perform further analysis (Fig. S2).

### CFTR is required for thermal stress-induced alteration

We next examined the contribution of epithelial Cl^−^ transport to thermal stress-induced alterations in R_TE_ and I_SC_ of MTE monolayers. Transcellular Cl^−^ transport via apical CFTR comprises majority of transcellular anion flux (Sheppard and Welsh 1999; Fuller and Benos 1992). At 37°C, Cl^−^-free bath solution or CFTRinh-172 was applied to block overall epithelial Cl^−^ transport or CFTR. Blockade of CFTR led to an increment in R_TE_ and a drop in I_SC_ (Fig. 3a and 3b, black line). However, Cl^−^-free bath solution caused a transient increment, followed by a slow decline to 0.24 ± 0.03 kΩ × cm^2^ in R_TE_ and almost zero in I_SC_ (Fig. S3). Therefore, we could not perform further analysis. During thermal stress, R_TE_ and I_SC_ were changed in a two-phase manner in the presence of CFTRinh-172, which was similar to the saline group (Fig. 3a and 3b, red line). CFTRinh-172 significantly increased thermal stress-sensitive R_TE_ for both phases (Fig. 3c, all *P* < 0.01). CFTRinh-172 significantly decreased thermal stress-sensitive I_SC_ for both phases (Fig. 3d and 3e, *P* < 0.05). These findings suggest that CFTR plays a role in changes in R_TE_ and I_SC_ caused by thermal stress.

### Steady-state elevated temperature increases transcellular ion channel activity

To confirm the two-phase alteration in I_SC_ following a gradual rise in bath temperature, we alternatively mounted MTE monolayers to Ussing chambers that were already set at 40°C. Interestingly, elevated and stable total I_SC_ levels were observed under this steady-state, elevated temperature condition, which was different from the two-phase manner when the temperature was gradually elevated (Fig. 4a, 4c, and 4e, basal I_SC_ of the black line and red line). These observations suggested that the time course of thermal stress determined the severity of impaired transcellular ion transport in MTE monolayers. To inhibit the activity of ion channels and transporters, including ENaCs, CFTR, and Na^+^/K^+^-ATPase, their specific inhibitors, including amiloride, CFTRinh-172, and ouabain, were added to a bath solution at 37°C or 40°C. I_SC_ was inhibited by these specific inhibitors at 37°C and 40°C (Fig. 4a, 4c, and 4e). We computed total I_SC_, specific inhibitor-resistant I_SC_, and the difference between these two I_SC_ (Figs. 4b, 4d, and 4f). The difference between total I_SC_ and specific inhibitor-resistant I_SC_ reflected the activity of corresponding ion channels or transporters, and we found that thermal stress increased the activity of ENaCs, CFTR and Na^+^/K^+^-ATPase (all *P* < 0.05). We also computed the basal R_TE_ value and post-inhibitor value applying at 37°C or 40°C (Fig. 4g). The mean basal R_TE_ level at 40°C (0.57 ± 0.05 kΩ × cm^2^) was significantly less than that at 37°C (1.08 ± 0.06 kΩ × cm^2^) (*P* < 0.001). An increase in R_TE_ was observed in the presence of amiloride and CFTRinh-172 (all *P* < 0.05), but not ouabain (all *P >* 0.05), at 37°C and 40°C. These data suggest that a steady-state elevated temperature increases the activity of ENaCs, CFTR, and Na^+^/K^+^-ATPase.

### Acrolein downregulates R_TE_ and I_SC_

Our previous studies showed that formaldehyde and crotonaldehyde regulated ENaCs in alveolar epithelial cells (Cui et al. 2016; Li et al. 2017). Acrolein is an industrial chemical of high toxicity and a toxic combustion product (Stevens and Maier 2008), and concentrations of which can vary widely in different fire and tobacco smoke. One kilogram of different types of wood can produce 0.374-2.35 mM of acrolein (Faroon et al. 2008). In the case of burning of tobacco, the acrolein concentrations are 34-503 μM when smoke from a single cigarette is bubbled through 10 ml of buffered saline (Burcham et al. 2010). To examine the effects of acrolein on the permeability of MTE monolayers, acrolein (approximately 500 μM) was applied to monolayers bathed at 37°C and at a steady-state of 40°C. A total of 500 μM acrolein decreased R_TE_ and I_SC_ at 37°C and 40°C (Fig. 5a and 5b). To compare the difference in effects of acrolein at 37°C and 40°C, the dose-response curve for acrolein was generated by fitting raw data points with the Hill equation. By fitting raw R_TE_ data, we found that the *K_i_* value for acrolein was 120.22 μM at 37°C and 79.43 μM at 40°C (Fig. 5c). By fitting raw I_SC_ data, we found that the *K*i value for acrolein was 125.89 μM at 37°C and 77.62 μM at 40°C (Fig. 5d). In HBE monolayers, a reduction of R_TE_ and I_SC_ caused by 500 μM acrolein was observed, which was similar to the result from MTE monolayers (Fig. 5e and 5f). In the R_TE_ dose-response curve of HBE monolayers, the *K*i value for acrolein was 83.18 μM at 37°C and 97.73 μM at 40°C (Fig. 5g). In the I_SC_ dose-response curve of HBE monolayers, the *K*i value for acrolein was 125.89 μM at 37°C and 112.21 μM at 40°C (Fig. 5h). Our data suggest that acrolein impairs airway epithelial permeability in an atypical concentration-dependent manner and an elevated temperature exacerbates this damaging process.

### Na^+^ transport participates in acrolein-induced reduction of R_TE_ and I_SC_

We examined the role of epithelial Na^+^ transport on the decrease in R_TE_ and I_SC_ by acrolein in MTE monolayers. At the basal level of 37°C, Na^+^-free bath solution, amiloride, or ouabain was used to block overall epithelial Na^+^ transport, ENaCs or Na^+^/K^+^-ATPase. After pre-inhibition of these Na^+^ transport pathways, 500 μM acrolein was added to the apical bath solution, and changes in R_TE_ and I_SC_ caused by acrolein were observed. At 37°C, Na^+^-free bath solution and amiloride, but not ouabain, caused an increment in R_TE_ (Fig. 6a, black line). Additionally, a drop in I_SC_ induced by Na^+^-free bath solution, amiloride, and ouabain, was observed (Fig. 6b, black line). During exposure of 500 μM acrolein, R_TE_ was decreased in the presence of Na^+^-free bath solution, amiloride, and ouabain (Fig. 6a, blue line). Acrolein continuously decreased I_SC_ in the presence of amiloride but showed no obvious effect in the presence of Na^+^-free bath solution and ouabain (Fig. 6b, blue line). We calculated acrolein-sensitive R_TE_(ΔR_TE_, Fig. 6c), and found that Na^+^-free bath solution, amiloride, and ouabain significantly increased acrolein-sensitive R_TE_ (*P* < 0.05). Acrolein-sensitive I_SC_(ΔI_SC_, Fig. 6d) was also computed, and it was significantly decreased by Na^+^-free bath solution, amiloride, and ouabain (all *P* < 0.01). These data suggest that changes in R_TE_ and I_SC_ induced by acrolein may be related to transcellular pathways that are mediated by ENaCs and Na^+^/K^+^-ATPase.

### CFTR mediates the acrolein-induced reduction in R_TE_ and I_SC_

We further examined the role of CFTR in the decrease in R_TE_ and I_SC_ by acrolein in MTE monolayers. At basal level of 37°C, CFTRinh-172 was used to inhibit CFTR, and then 500 μM acrolein was added to the monolayers. Changes in R_TE_ and I_SC_ caused by acrolein were observed. At the basal level, CFTRinh-172 increased R_TE_ (Fig. 7a, black line) and decreased I_SC_ (Fig. 7b, black line). During exposure of acrolein, R_TE_ and I_SC_ were decreased (Figs. 7a and 7b, blue line). CFTRinh-172 significantly increased acrolein-sensitive R_TE_ (Fig. 7c, *P* < 0.01) and markedly decreased acrolein-sensitive I_SC_ (Fig. 7d, *P* < 0.001). These findings suggest that CFTR may participate in the reduction of R_TE_ and I_SC_ caused by acrolein.

### Thermal stress and acrolein impair tight junctions

Our findings suggested that the transcellular pathway is required for the effects of thermal stress and acrolein on permeability of MTE monolayers. Therefore, we next examined if thermal stress and acrolein affect the paracellular pathway. We observed the structure and distribution of the tight junction protein ZO-1 in MTE monolayers. In MTE monolayers, ZO-1 is one of the most widely studied scaffold proteins and is uniformly expressed after an air-liquid interface is achieved (Umeda et al. 2006). Confocal microscopy showed that thermal stress disrupted the integrity of ZO-1 (Fig. 8e) compared with the control group (Fig. 8b). Damage to the structure of ZO-1 was also observed in acrolein-treated monolayers (Fig. 8h). Moreover, synergistic treatment of thermal stress and acrolein further exacerbated the damage to ZO-1 (Fig. 8k) compared with thermal stress (Fig. 8e) or acrolein group (Fig. 8h). We quantified fluorescence intensity and it was consistent with that from the representative confocal images (Fig. 8m). To determine the effects of thermal stress and acrolein on cell detachment and death of MTE monolayers, we tested cell number and viability of monolayers post-thermal stress and after acrolein. We found no significant cell detachment or death under these conditions (Fig. S4, all *P* > 0.05). We examined expression of occludin and claudins, but their expression levels were insufficient for visualization (Fig. S6). The FITC-dextran assay was performed to further determine the effects of thermal stress and acrolein on paracellular permeability of MTE monolayers. We found that thermal stress and acrolein significantly increased the permeability of MTE monolayers to FITC-dextran (Fig. S6p, *P* < 0.001).

## Discussion

Smoke inhalation injury is the main cause of death during a fire, and it is an important risk factor for increasing morbidity and mortality (You et al. 2014). Pathogenic factors in inhalation injury, such as heat and toxic chemicals, cause direct injury to the airway epithelial barrier. In this study, we aimed to investigate the effects of thermal stress and acrolein on permeability of the airway epithelial barrier. MTE cells were isolated from mouse trachea and used to culture the model of the air-liquid interface (MTE monolayers). This *in vitro* model mainly contained goblet and cilia cells and imitated the physiological state of the normal airway epithelium (Hou et al. 2019). MTE monolayers are suitable for studying respiratory diseases, and they have been applied in toxicology, respiratory tract infection, ion transport, cell carcinogenesis, and other experiments (Davidson et al. 2000; Horani et al. 2013). In our study, R_TE_ and I_SC_ were markedly changed in thermal stress or acrolein-treated MTE monolayers. The changes in R_TE_ and I_SC_ caused by thermal stress or acrolein were related to the transcellular pathway, which was mediated by ENaCs, CFTR, and Na^+^/K^+^-ATPase. Additionally, thermal stress or acrolein regulated the paracellular pathway, which was mediated by tight junctions. These findings suggest the potential roles of the transcellular and paracellular pathways in thermal stress or acrolein in damaging the airway epithelial barrier. Our findings may provide new directions for the understanding, diagnosis, and treatment of smoke inhalation injury.

Transcellular and paracellular pathways are two essential determinants of R_TE_ and I_SC_. In the current study, during thermal stress, R_TE_ and I_SC_ of MTE monolayers were changed in a two-phase (P1 and P2) manner. I_SC_ was increased at P1, which could have been due to enhanced ion transport by thermal stress. Concurrently, increased permeability led to a reduction in R_TE_. The reverse change in R_TE_ and I_SC_ indicated that monolayers at P1 still had normal physiological characteristics. Subsequently, R_TE_ and I_SC_ were decreased at P2. These findings could have resulted from damaged gap and tight junctions, which were shown by confocal images. Another explanation for our findings is ion diffusion across MTE monolayers upon depletion of ATP. Therefore, we speculate that effects caused by thermal stress are reversible at P1 and irreversible at P2. A previous study provided the basis for our speculation (Tobey et al. 1999) in which increased I_SC_ and decreased R_TE_ were observed during application of heat to the esophageal epithelium, and recovery was observed when the temperature returned to 37°C. Cell death and detachment cause increased permeability of monolayers. However, our data from cell counting and viability ruled out this possibility. To overcome the difference between species, we cultured HBE monolayers and monitored R_TE_ and I_SC_. We found that R_TE_ of HBE monolayers was decreased by thermal stress, which was similar to the result from MTE monolayers. In contrast, thermal stress increased I_SC_ of HBE monolayers to a high and stable level, which was similar to P1 in MTE monolayers. However, we could not monitor the decreased trace of I_SC_ (i.e., P2 in MTE monolayers). These findings indicated that HBE monolayers showed lower thermal sensitivity and higher thermal resistance than primary MTE monolayers.

Various time courses of thermal stress caused obvious different regulation in I_SC_ in our study. We observed an elevated I_SC_ trace at P1 and a declined trace at P2 when the temperature was gradually raised to 40°C from 37°C. However, under the condition of a steady-state at 40°C, an elevated and stable I_SC_ trace was visualized instead of two phases. In a previous study, researchers found elevated stable I_SC_ levels of rabbit esophageal epithelium in an Ussing chamber system in which the temperature had already been set to 49°C (Tobey et al. 1999). These divergent responses in I_SC_ and R_TE_ to different time courses of thermal stress could be due to the response of the individual transport system to thermal stress. Host cell adaption to environmental stress could explain the diverse observations between the two procedures of thermal stress.

The paracellular and transcellular pathways are vital for establishing or dissipating ion concentration gradients and thus are important for determining the ionic composition of the apical compartment and net volume flow (Flynn et al. 2009). Therefore, both pathways work in concert and are functionally matched to meet the transport requirements of the specific tissue. The paracellular pathway is mainly formed by tight junctions, and this pathway is located near the apical side of the cells and the lateral intercellular space (Tsukita and Furuse 2002; Van Itallie and Anderson 2006). Tight junctions form the functional and structural boundary that separates apical and basolateral compartments and also determine the ion transport properties of the paracellular pathway. Tight junctions are composed of a complex of proteins that determine ion selectivity and conductance of the paracellular pathway (Farquhar and Palade 1963). ZO-1 is one of the most widely studied tight junction protein (Umeda et al. 2006) and is uniformly expressed after the air-liquid interface is achieved (Kuroishi et al. 2009). We found that thermal stress or acrolein decreased ZO-1 expression and disrupted its tight structure, which indicated that thermal stress or acrolein regulated the paracellular ion transport pathway. Inhibitors (e.g., Amiloride, ouabain and CFTRinh-172) can only block ion channels and transporters that belong to the transcellular pathway and inhibit the function of ion transport. Therefore, inhibitors for ENaCs, CFTR, and Na^+^/K^+^-ATPase might not significantly affect these proteins. In our study, the FITC-dextran flux assay showed increased FITC-dextran permeability in thermal stress or acrolein-treated MTE monolayers. Thermal stress or acrolein not only damaged the complete structure of the tight junction but also increased its permeability.

In the current study, when adding acrolein to an apical bath solution, the bath solution was blown up to 10 times to ensure even distribution of acrolein instead of gas bubbling. Maintaining bubbling of a bath solution facilitates loss of acrolein. This situation was supported by our results for comparing two experimental conditions with and without air bubbling (Fig. S5). A total of 500 μM acrolein adversely affected the integrity of the airway epithelial barrier, which was supported by a reduction in R_TE_ and I_SC_, a disrupted ZO-1 structure, as shown by confocal images, and the increased FITC-dextran permeability in MTE monolayers. Therefore, 500 μM acrolein caused irreversible damage to the airway epithelial barrier. Through a fitted R_TE_/I_SC_ dose-response curve for acrolein in MTE monolayers, we found that the *K*i value for acrolein was lower at 40°C than at 37°C. Therefore, the same dose of acrolein caused more serious damage to R_TE_/I_SC_ under thermal stress conditions compared with 37°C. Moreover, the damage to the integrity and permeability of tight junctions caused by acrolein was more severe at 40°C than at 37°C. The synergistic effects of thermal stress and acrolein induced further impairment to the airway epithelial barrier. Our previous studies indicated that oxidative aldehydes, including formaldehyde and crotonaldehyde, inhibited activity of ENaCs by causing activation of reactive oxygen species (ROS) in alveolar epithelium cell monolayers (Cui et al. 2016; Li et al. 2017). Modulation of antioxidant enzymes affected the thermal sensitivity of respiratory cells. Additionally, lowering superoxide dismutase enzyme levels resulted in a significant reduction in thermal resistance (Omar et al. 1987). However, overexpression of manganese superoxide dismutase by stable transfection provided cellular resistance against the cytotoxic effect of hyperthermia (Li and Oberley 1997; Kuninaka et al. 2000). Therefore, the synergistic effects of thermal stress and acrolein in the present study may have resulted from increased thermal sensitivity to cells, which was caused by acrolein-activated oxidative signaling.

As mentioned above, damage to the airway epithelium due to acrolein mainly results from its downstream signal via cellular oxidative stress, such as glutathione depletion, and subsequently ROS stimulation (Wang et al. 2009). In addition to ENaCs, Na^+^/K^+^-ATPase, and CFTR, some other proteins, including KCs, the Ca^2+^-activated Cl^−^ channels (CaCCs), SCL26A9, and Na^+^/K^+^/2Cl^−^ (NKCC), play roles in ion transport in the respiratory epithelium. Additionally, these proteins may be regulated by acrolein by induction of oxidative stress (Londino et al. 2017). As the major Ca^2+^-activated KCs of ciliated cells, K_Ca_1.1/KCNMA1 plays a role in maintaining fluid secretion and is affected by oxidative stress (Manzanares et al. 2011; Kis et al. 2016; Hermann et al. 2015). Moreover, exposure to tobacco smoke inhibits K_Ca_1.1/KCNMA1 and results in a reduction in airway surface liquid in human bronchial epithelial cells (Sailland et al. 2017). Anoctamin 6 functions as a CaCCs, and a Ca^2+^-dependent phospholipid scramblase, which are stimulated by an increase in ROS and subsequent peroxidation of membrane lipids (Schreiber and Ousingsawat 2018; Scudieri et al. 2015). NKCC is located on the basolateral membrane of the tracheal epithelium and is an important conduit for Cl^−^ entry in the liquid-transporting epithelium (Gillie et al. 2001). A previous study reported that NKCC activation may contribute to the protective system against ROS-mediated damage to the airway epithelium (Matsuno et al. 2008). These studies may provide proof of acrolein damaging KCs, CaCCs, and NKCC in the airway epithelium.

In addition to the effects of smoke inhalation on the tracheal epithelium, we also found interactions between different ion transport systems. Apical and basolateral ion transport systems play diverse roles in the relationship between R_TE_ and I_SC_. In our study, blockade of Na^+^ entry or Cl^−^ secretion via apical ENaCs and CFTR, respectively, caused a reciprocal change in R_TE_ and I_SC_. In contrast, inhibition of basolateral Na^+^/K^+^-ATPase resulted in a reduction in R_TE_ and I_SC_. Additionally, a Cl^−^-free solution showed a similar effect as ouabain in inhibiting I_SC_. This finding indicates the potential dependence of Na^+^/K^+^-ATPase on anion transport or interactions between vertical Cl^−^ and K^+^ transporters. This interesting similar effect can be excluded for the associations between Cl^−^ and Na^+^ transporters, because a Na^+^-free solution led to an opposite effect on R_TE_ and I_SC_. Our study showed that the apical ENaCs and CFTR modulated R_TE_.

There are some limitations to this study, which need to be addressed in the future. Although the MTE *in vitro* model imitates the physiological state and complies with the morphological characteristics of the normal airway epithelium, there are still some differences from the *in vivo* environment. However, an *in vivo* model does not allow us to precisely measure R_TE_ and I_SC_. Future studies need to focus on the effects of thermal stress or acrolein on inflammatory signal transduction pathways at the molecular level. In addition to the oxidative stress pathway, the mitogen-activated protein kinase (MAPK) pathway appears to play a role in regulating the airway epithelial barrier (Huang et al. 2016; Lee et al. 2018). A previous study suggested that MAPK Hog1 phosphorylation was activated to a peak at approximately 5 min after heat shock, and phosphorylated Hog1 declined to the basal level at 30 min (Dunayevich et al. 2018). Researchers reached a similar conclusion in another study, where they found that heat-activated MAPK signaling was elevated to a peak at 5 min, and activation was weakened or even disappeared over time (Dong et al. 2015). In the present study, application of thermal stress to MTE monolayers caused a bi-phasic change, which indicated a sophisticated condition and a long time course. An *in vivo* model can show higher sensitivity and resistance to thermal stress than an *in vitro* model. Therefore, we ultimately to perform an *in vivo* study of thermal stress or acrolein-induced MAPK signaling.

Thermal stress and acrolein are the two main pathogenic factors of smoke inhalation injury. Our study suggests that these two factors damage the integrity of the airway epithelial barrier. The underlying mechanisms are related to the transcelluar pathway (mediated by ion channels and transporters) and the paracellular pathway (mediated by tight junctions). Morever, to the best of our knowledge, we have shown for the first time the synergistic effects of thermal stress and acrolein on impairment of the airway epithelial barrier.

## Authors’ contributions

Hong-Long Ji conceived study, designed experiments, edited manuscript, and approved submission. Jianjun Chang, Zaixing Chen, and Hong-Guang Nie performed experiments, analyzed data, and plotted graphs. Jianjun Chang, Runzhen Zhao, Hong-Guang Nie, and Zaixing Chen prepared manuscript.

## Abbreviations

ENaCs: epithelial Na^+^ channels
CFTR: cystic fibrosis transmembrane regulator
R_TE_: transepithelial resistance
I_SC_: short-circuit current
MTE: mouse tracheal epithelial
FITC: fluorescein isothiocyanate
HBE: human bronchial epithelial
DMSO: dimethyl sulfoxide
ZO-1: zonula occludens-1
P1: phase 1
P2: phase 2
ASI: amiloride-sensitive I_SC_
KCs: K^+^ channels
ROS: reactive oxygen species
CaCCs: Ca^2+^-activated Cl^−^ channels
NKCC: Na^+^/K^+^/2Cl^−^
MAPK: mitogen-activated protein kinase

## Acknowledgments

This work was supported by the grants from the National Institute of Health (NIH HL134828), and the National Natural Science Foundation of China (NSFC 81670010).

## Figure Legends

**Fig. 1 Thermal stress alternates bioelectric features in mouse tracheal epithelial (MTE) and human bronchial epithelial (HBE) monolayers. (a)** Representative transepithelial resistance (R_TE_) trace in MTE monolayer mounted on an Ussing chamber setup. R_TE_ was decreased in a two-phase manner (P1 and P2 were labeled) during the temperature of the bath solution was raised to 40°C (red line) from 37°C (black line). When the temperature rose to 38.5°C, the changes in R_TE_ and I_SC_ were observed, and it took 4.7 min for the course from 37°C to 40°C. **(b)** Short-circuit current (I_SC_) trace recorded simultaneously in the same MTE monolayer. P1 and P2 of I_SC_ were defined using the R_TE_ time course. I_SC_ was increased at P1 and decreased at P2. **(c)** Average R_TE_ levels of MTE monolayers at different time points during thermal stress. Student’s t-test. \*\*\**P* < 0.001. *n* = 20. **(d)** Average I_SC_ levels of MTE monolayers. Student’s t-test. \*\*\**P* < 0.001. *n* = 20. **(e)** Time for reducing half of the total R_TE_ (τ_1/2_). The τ_1/2_ value was computed by fitting R_TE_ raw data with the ExpDec1 function. (*y*_0_ = 0.36 ± 0.04, *A* = 0.87 ± 0.04 [P1]; *y*_0_ = −0.87 ± 0.11, *A* = 3.22 ± 0.36 [P2]). Student’s t-test. \*\*\**P* < 0.001. *n* = 12. **(f)** Representative R_TE_ trace in HBE monolayer. R_TE_ was decreased during the temperature of the bath solution was elevated to 40°C (red line) from 37°C (black line). **(g)** I_SC_ trace recorded simultaneously in the same HBE monolayer. I_SC_ was increased to a high and stable level during thermal stress. **(h)** Average R_TE_ levels of HBE monolayers under the conditions of 37°C and 40°C. Student’s t-test. NS, no significance. *n* = 6. **(i)** Average I_SC_ levels of HBE monolayers. Student’s t-test. \**P* < 0.05. *n* = 6.

**Fig. 2 Na^+^ transport mediates thermal stress-induced bioelectric changes in MTE monolayers. (a, b)** Representative R_TE_ and I_SC_ traces in the presence of saline bath solution (control group, solid line), Na^+^-free bath solution (dashed lines), amiloride (Amil, 100 μM, dotted lines), or ouabain (1 mM, dashed-dotted lines). At 37°C (black lines), Na^+^-free bath solution, amiloride or ouabain was applied at the time pointed by the arrow. And the temperature of the bath solution was elevated to 40°C (red lines). When the temperature rose to 38.5°C, the changes in R_TE_ and I_SC_ were observed. **(c)** Average thermal stress-sensitive R_TE_ levels (ΔR_TE_, the difference between the initial R_TE_ and ending R_TE_ within one phase [P1 or P2]). Mann-Whitney U test and Student’s t-test. \*\**P* < 0.01. NS, no significance. *n* = 27. **(d)** Average thermal stress-sensitive R_TE_ levels (ΔR_TE_, the difference between the initial R_TE_ and ending R_TE_ during the entire process of thermal stress). Student’s t-test. \**P* < 0.05 and \*\**P* < 0.01. *n* = 17. **(e)** Average thermal stress-sensitive I_SC_ levels at P1 (ΔI_SC_, the difference between the basal I_SC_ and the peak I_SC_ at P1). Student’s t-test. \**P* < 0.05 and \*\*\**P* < 0.001. *n* = 29. **(f)** Average thermal stress-sensitive I_SC_ levels at P2 (ΔI_SC_, the difference between the peak I_SC_ and the ending I_SC_ at P2). Student’s t-test. \*\*\**P* < 0.001. *n* = 20.

**Fig. 3 CFTR is involved in thermal stress-induced bioelectric changes in MTE monolayers (a, b)** Representative R_TE_ and I_SC_ traces in the presence of saline bath solution (control group, solid line) and CFTRinh-172 (CFTRinh, 20 μM, dashed line). Segments of traces recorded at different bath temperatures were shown as black (37°C) and red lines (40°C). Arrows indicated the time to add CFTRinh-172. When the temperature rose to 38.5°C, the obvious bioelectric changes were observed. **(c)** Average thermal stress-sensitive R_TE_ levels (ΔR_TE_, the difference between the initial R_TE_ and ending R_TE_ within one phase [P1 or P2]). Student’s t-test. \*\**P* < 0.01 and \*\*\**P* < 0.001. *n* = 18. **(d)** Average thermal stress-sensitive I_SC_ levels at P1 (ΔI_SC_, the difference between the basal I_SC_ and the peak I_SC_ at P1). Student’s t-test. \**P* < 0.05. *n* = 9. **(e)** Average thermal stress-sensitive I_SC_ levels at P2 (ΔI_SC_, the difference between the peak I_SC_ and the ending I_SC_ at P2). Student’s t-test. \**P* < 0.05. *n* = 9.

**Fig. 4. Activation of ENaCs, CFTR, and Na^+^-K^+^-ATPase of MTE monolayers by thermal stress. (a, c, and e)** Representative I_SC_ traces before and after the addition of amiloride (100 μM), CFTRinh (20 μM), or ouabain (1 mM) at 37°C (black lines) and 40°C (red lines). Arrows indicated the time to add inhibitors. **(b, d and f)** Average I_SC_ levels. Total I_SC_ meant the basal I_SC_ level; +Amil I_SC_, +CFTRinh I_SC_, and +Ouabain I_SC_ meant inhibitor-resistant I_SC_ level; ASI (amiloride-sensitive I_SC_), CFTRinh I_SC_ and Ouabain I_SC_ respectively reflected the activity of each ion channel and was calculated by the difference between the basal I_SC_ and inhibitor-resistant I_SC_. Student’s t-test and Mann-Whitney U test. \**P* < 0.05 and \*\**P* < 0.01. NS, no significance. *n* = 18. **(g)** Average R_TE_ levels. Student’s t-test and Mann-Whitney U test. \**P* < 0.05 and \*\*\**P* < 0.001. NS, no significance. *n* = 36.

**Fig. 5 Acrolein impairs bioelectric features in MTE and HBE monolayers. (a, b)** Representative R_TE_ and I_SC_ traces in the presence of 5, 50, and 500 μM acrolein at 37°C (black lines) and 40°C (red lines) in MTE monolayers. Acrolein (ACR, 5, 50, and 500 μM) was pipetted to the bath as indicated by arrows. 500 μM acrolein decreased R_TE_ and I_SC_. **(c, d)** Dose-response curve for acrolein in MTE monolayers. Normalized R_TE_/I_SC_ points were fitted with the Hill equation (*START* = 1, *END* = 0, *n* = 22). The *K_i_* value of R_TE_ for acrolein was 120.22 μM at 37°C and 79.43 μM at 40°C; The *K*i value of I_SC_ for acrolein was 125.89 μM at 37°C and 77.62 μM at 40°C. **(e, f)** Representative R_TE_ and I_SC_ traces of HBE monolayers in the presence of 5, 50, and 500 μM acrolein under conditions of 37°C (black line) and 40°C (red line). Arrows indicated the time to add acrolein (ACR, 5, 50, and 500 μM). R_TE_ and I_SC_ were decreased by 500 μM acrolein. **(g, h)** Dose-response curve for acrolein in HBE monolayers. Normalized R_TE_/I_SC_ points were fitted by the Hill equation (*START* = 1, *END* = 0, *n* = 2). The *K*i value of R_TE_ for acrolein was 83.18 μM at 37°C and 97.73 μM at 40°C, respectively; the *K*i value of I_SC_ for acrolein was 125.89 μM at 37°C and 112.21 μM at 40°C, respectively.

**Fig. 6 Na^+^ transport is involved in acrolein-impaired bioelectric features in MTE monolayers. (a, b)** Representative R_TE_ and I_SC_ traces in the presence of saline bath solution (control group, solid lines), Na^+^-free bath solution (dashed lines), amiloride (Amil, 100 μM, dotted lines), or ouabain (1 mM, dashed-dotted line). At basal level (black lines), Na^+^-free bath solution, amiloride or ouabain was applied at the time pointed by the arrow. And then 500 μM acrolein (ACR, blue lines) was added. **(c)** Average acrolein-sensitive R_TE_ levels (ΔR_TE_, the difference between basal R_TE_ and acrolein-resistant R_TE_). Mann-Whitney U test. \**P* < 0.05. *n* = 16. **(d)** Average acrolein-sensitive I_SC_ levels (ΔI_SC_, the difference between basal I_SC_ and acrolein-resistant I_SC_). Student’s t-test. \*\**P* < 0.01 and \*\*\**P* < 0.001. *n* = 16.

**Fig. 7 CFTR mediates acrolein-impaired bioelectric features in MTE monolayers. (a, b)** Representative R_TE_ and I_SC_ traces in the presence of saline bath solution (control group, solid lines) and CFTRinh-172 (20 μM, dashed lines). Arrows showed the time for adding CFTRinh-172. Then monolayers were exposed to 500 μM acrolein (ACR, blue lines). **(c)** Average acrolein-sensitive R_TE_ levels (ΔR_TE_, the difference between basal R_TE_ and acrolein-resistant R_TE_). Student’s t-test. \*\**P* < 0.01. *n* = 11. **(d)** Average acrolein-sensitive I_SC_ levels (ΔI_SC_, the difference between basal I_SC_ and acrolein-resistant I_SC_). Student’s t-test. \*\*\**P* < 0.001. *n* = 11.

**Fig. 8 The synergistic effects of thermal stress and acrolein on ZO-1 tight junction in MTE monolayers.** Laser scanning confocol imaging for ZO-1 protein in MTE monolayers treated by 37°C **(a-c)**, 40°C for 120 min **(d-f)**, 500 μM acrolein **(g-i)**, and combination of 40°C and 500 μM acrolein **(j-1)**. Nucleus was labeled with hoechst (blue), and tight junction was labeled with ZO-1 (green). Scale bar: 50 μm. **(m)** Quantification of ZO-1 fluorescence intensity. Student’s t-test. \**P* < 0.05, \*\*\**P* < 0.001, &*P* < 0.05 and #*P* < 0.05. *n* = 12.

## Supplementary materals

### Supplemental Figure Legends

**Fig. S1.**
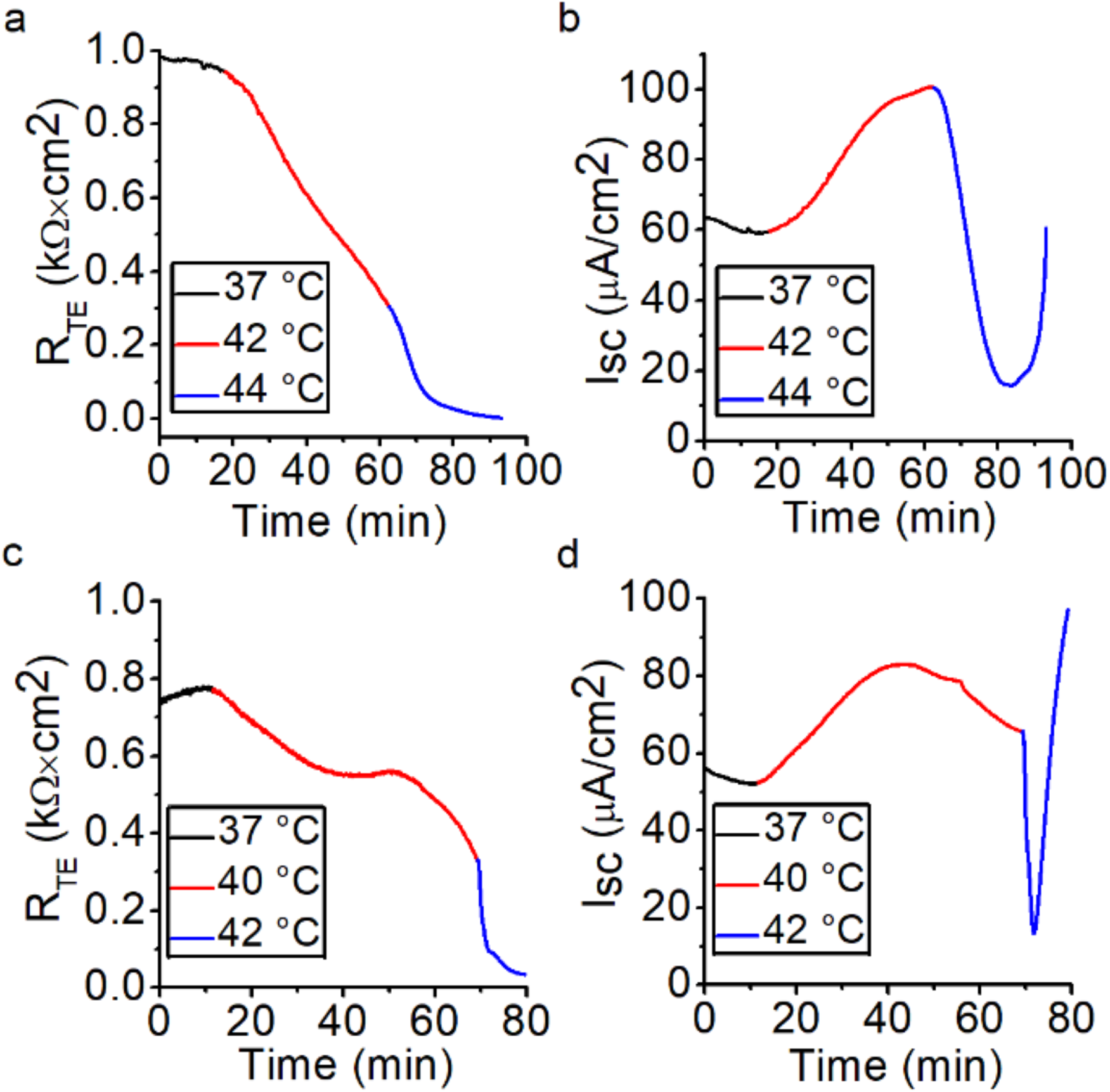
Extreme thermal stress disrupts bioelectric features in minutes in mouse tracheal epithelial (MTE) monolayers. **(a)** Representative transepithelial resistance (R_TE_) trace at 37°C, 42°C, and 44°C. R_TE_ was eliminated to zero at 44°C. **(b)** Short-circuit current (I_SC_) trace recorded simultaneously of the same monolayer. At 44°C, I_SC_ was decreased followed by a jump, probably because of the damage to tight monolayer. **(c)** Representative R_TE_ trace at 37°C, 40°C, and 42°C. R_TE_ was eliminated to zero at 42°C. **(d)** I_SC_ trace of the same monolayer. I_SC_ was decreased at 42°C and then shot up and out of the scale in minutes probably due to impaired monolayer.

**Fig. S2.**
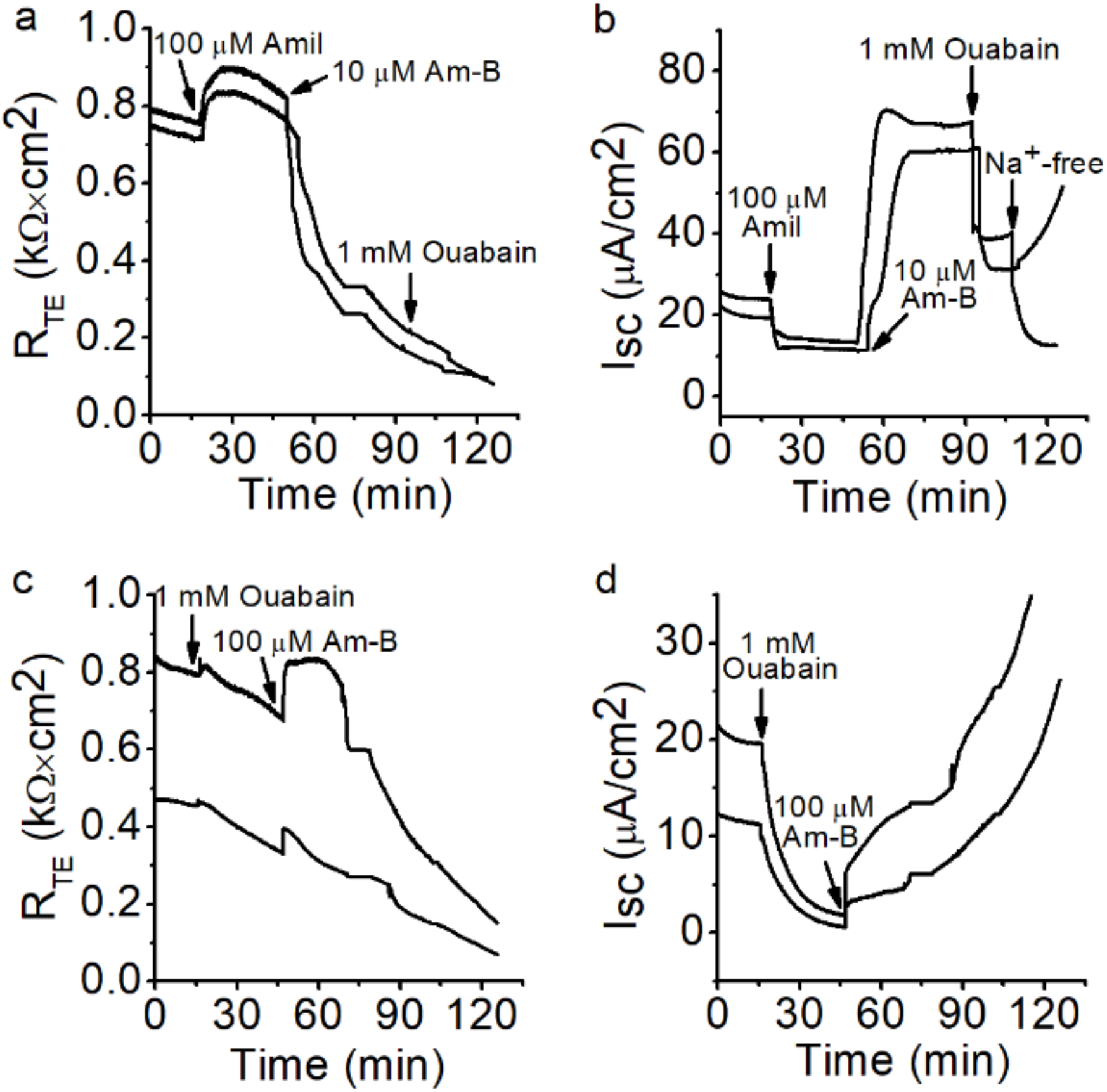
Long process of apical and basolateral-permeabilizing destabilizes R_TE_ and I_SC_ measurements in MTE monolayers. **(a)** Representative R_TE_ traces of apical-permeabilized MTE monolayers. After long time of apical permeabilizing process, R_TE_ was decreased near zero. **(b)** I_SC_ traces monitored synchronously of the same monolayers. I_SC_ was reduced followed by a jump. **(c)** Representative R_TE_ traces of basolateral-permeabilized MTE monolayers. R_TE_ was almost decreased to zero. **(d)** I_SC_ traces of the same monolayers. I_SC_ was decreased and then out of the scale upwardly probably due to the damage to tight monolayers.

**Fig. S3.**
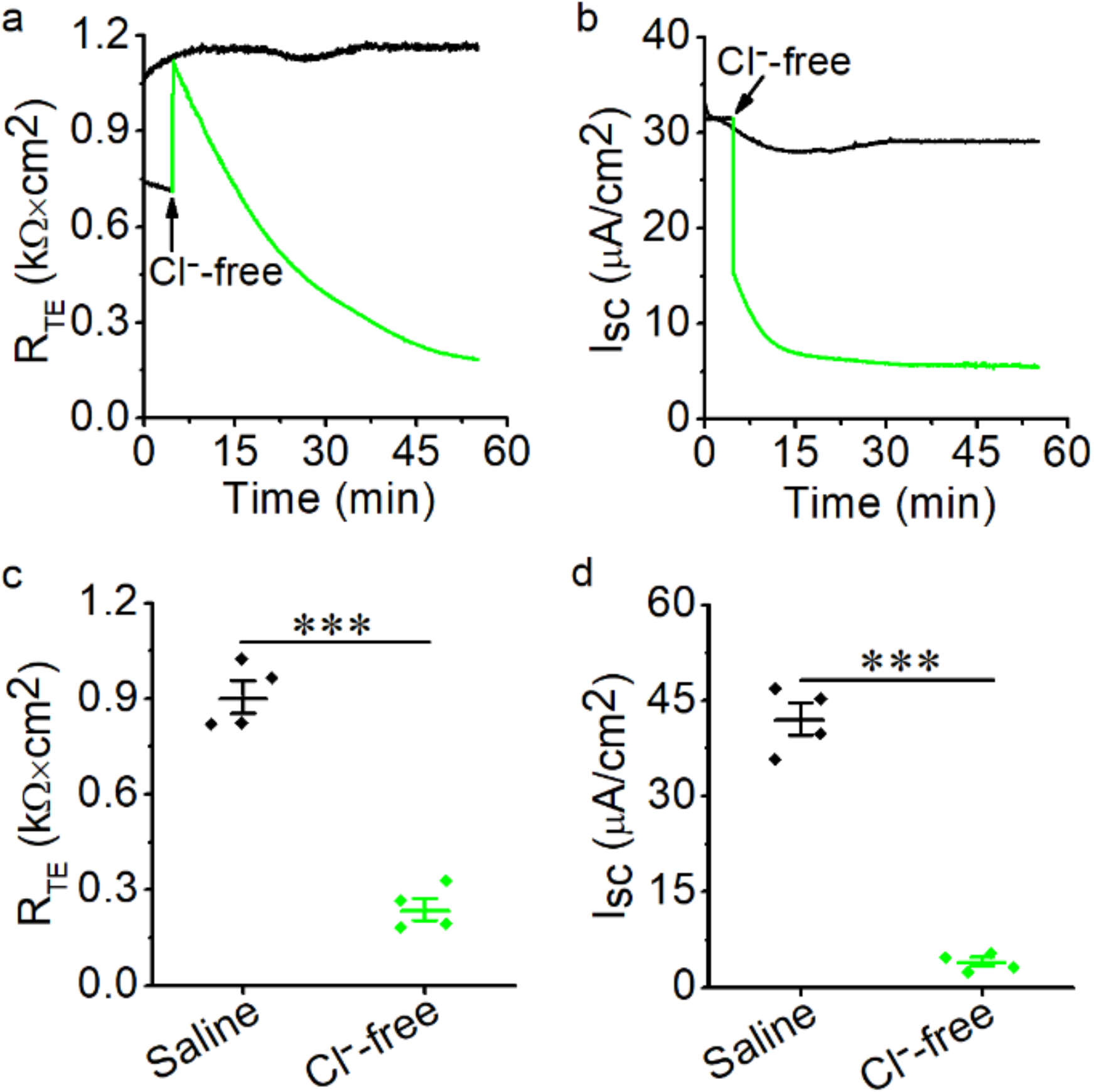
Cl^−^-free bath solution eliminates R_TE_ and I_SC_ levels in MTE monolayers. **(a)** Representative R_TE_ trace in the presence of saline bath solution (control group, black line) and Cl^−^-free bath solution (green line). Cl^−^-free bath solution caused the transient increment followed by the slow decline of R_TE_. **(b)** I_SC_ trace recorded synchronously of the same monolayer. **(c)** Average R_TE_ levels. R_TE_ was eliminated to 0.24 ± 0.03 kΩ × cm^2^ compared with the control group (0.91 ± 0.05 kΩ × cm^2^), probably due to impaired tight monolayers. Student’s t-test. *\*\*\*P* < 0.001. *n* = 8. **(d)** Average I_SC_ levels. Student’s t-test. *\*\*\*P* < 0.001. *n* = 8.

**Fig. S4.**
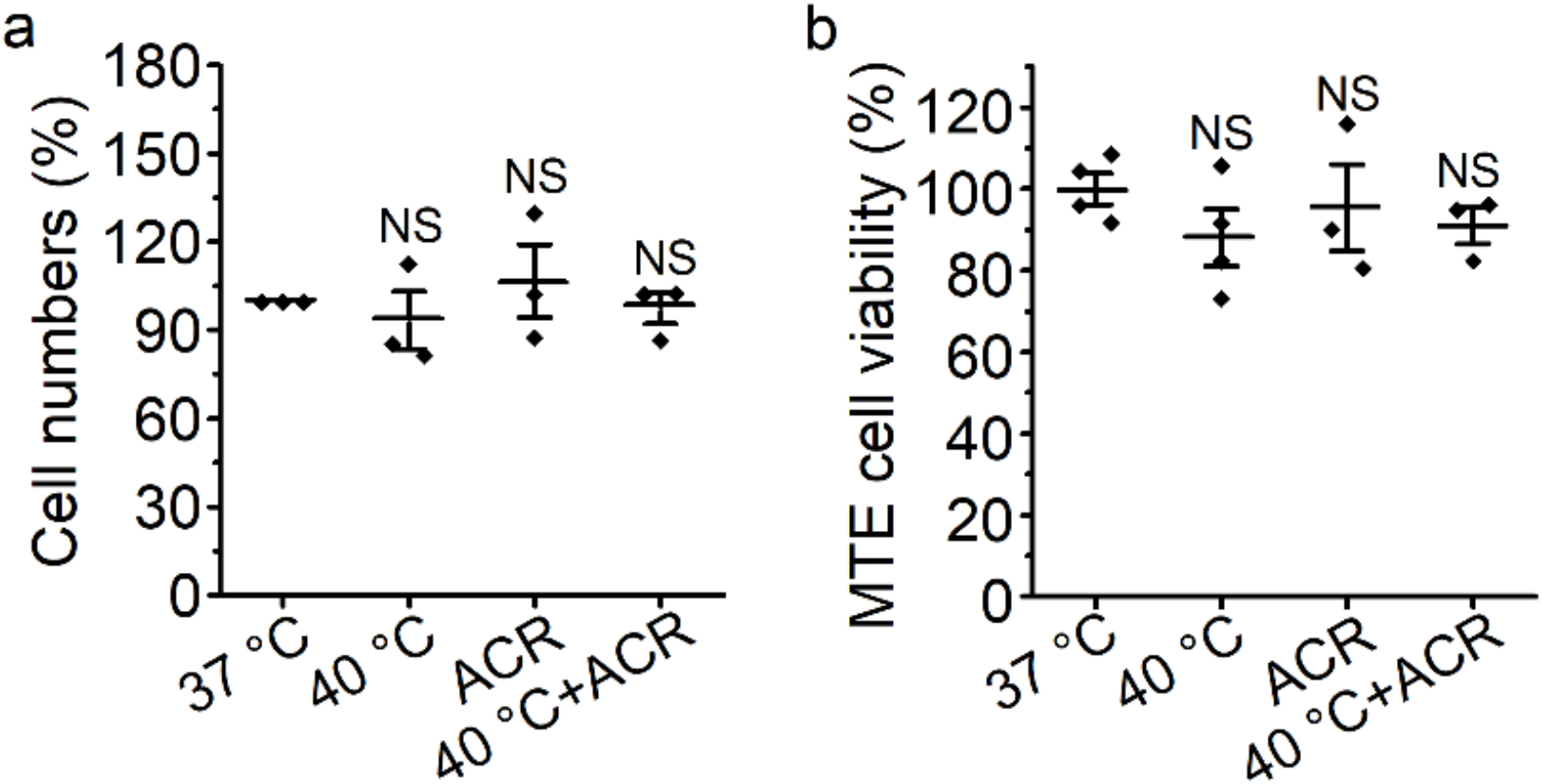
Cell count and viability in MTE monolayers. **(a)** Cell number of monolayers post the exposure of thermal stress (120 min) and acrolein (500 μM). Ten randomly selected images across the monolayers from at least 3 independent experiments were captured and counted for total cells with DAPI staining using a cell count plug-in of the ImageJ. No significant difference was observed among each group. Thermal stress and acrolein could not induce significant cell detachment of monolayers. NS, no significance. *n* = 12. **(b)** Cell viability of monolayers treated by thermal stress (120 min) and acrolein (500 μM). At the end of treatment, cell viability was measured with the cell counting kit-8 assay. Thermal stress and acrolein failed to cause obvious cell death of monolayers. NS, no significance. *n* = 12.

**Fig. S5.**
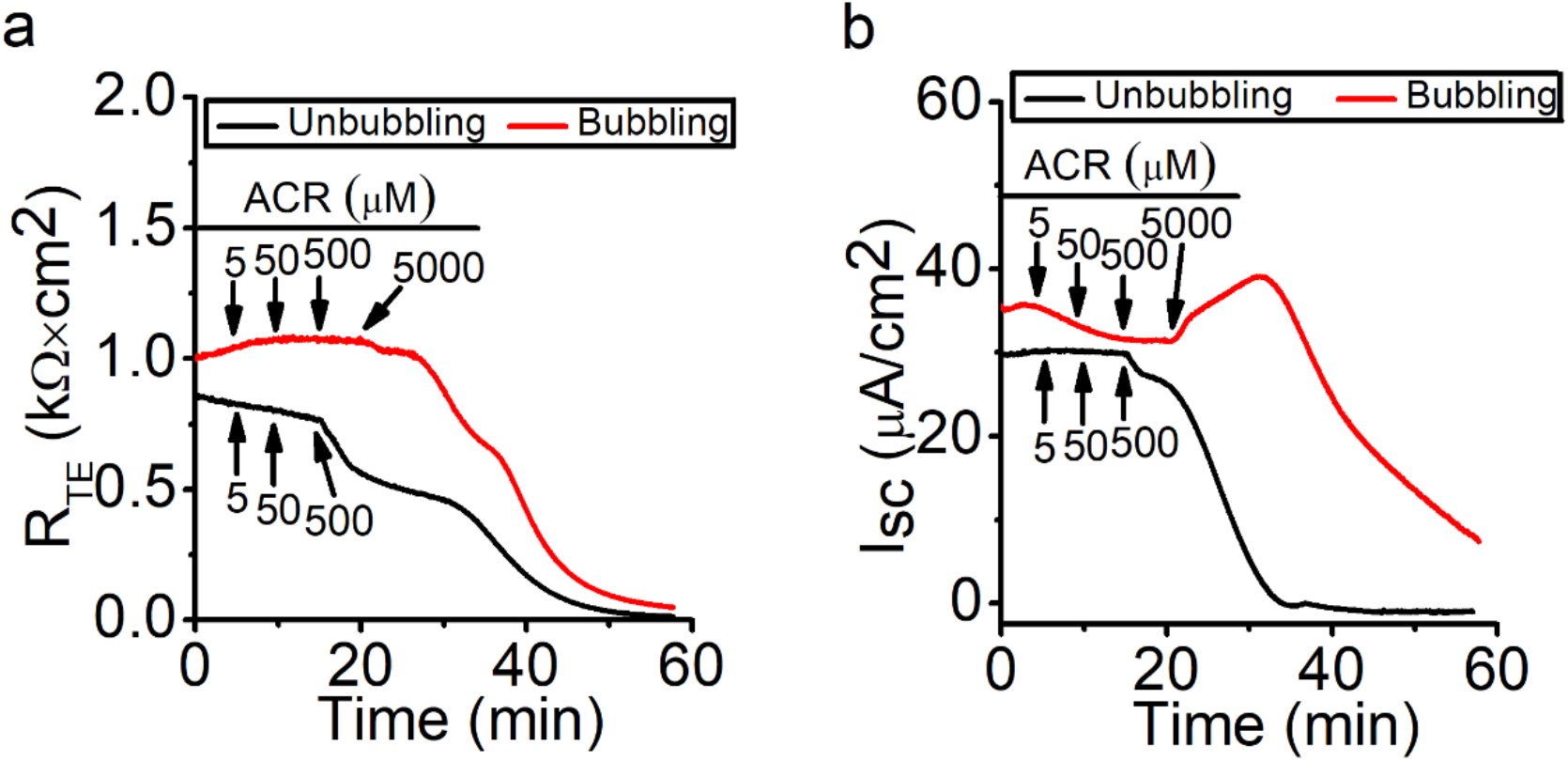
Comparison of the effects of acrolein on I_SC_ and R_TE_ in bubbled and unbubbled MTE monolayers. **(a)** Representative R_TE_ trace when unbubbling (black line) or bubbling (red line). **(b)** I_SC_ trace recorded simultaneously of the same monolayers. Keeping bubbling the bath solution facilitated the loss of acrolein and mitigated the effects of acrolein on R_TE_ and I_SC_.

**Fig. S6.**
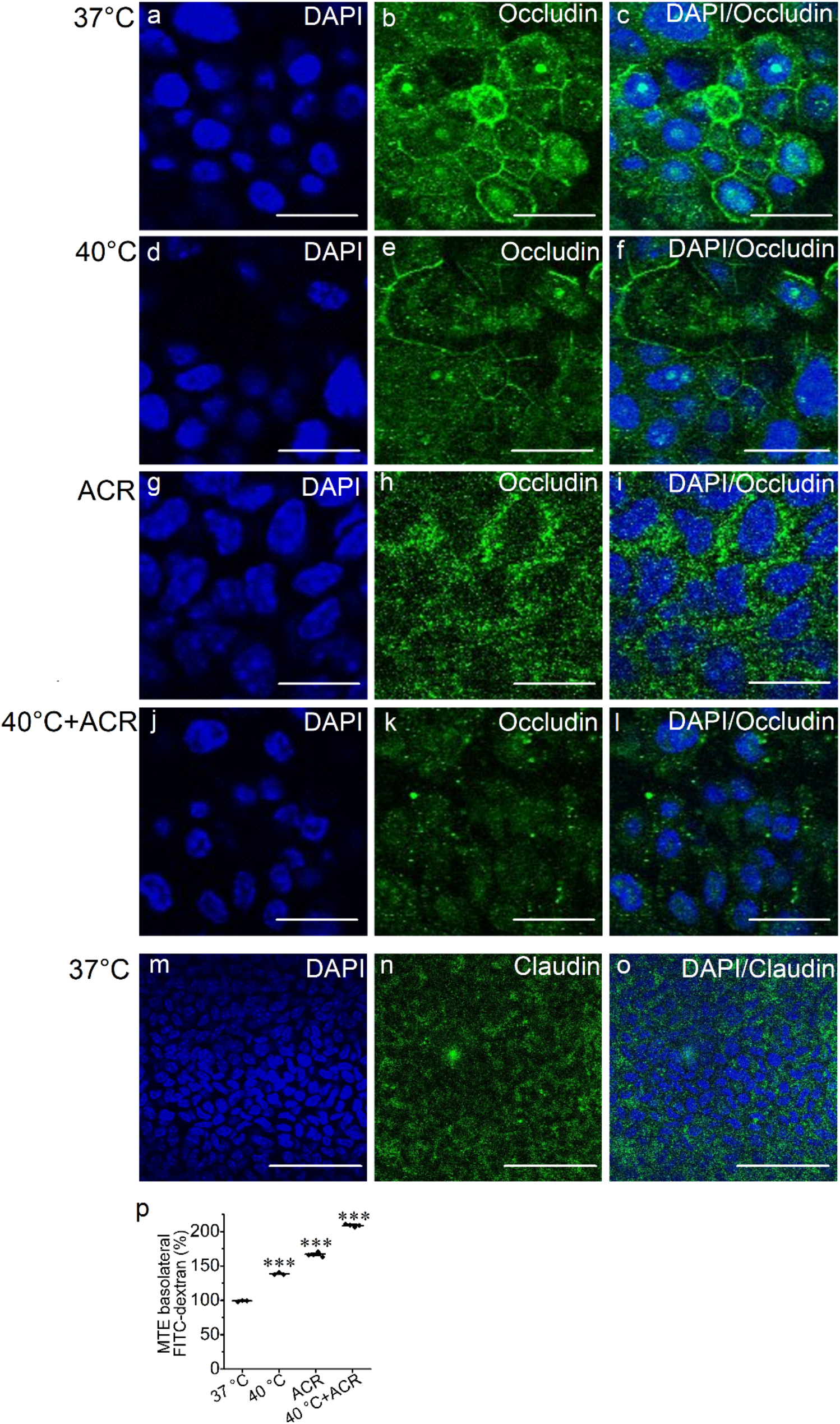
Thermal stress and acrolein impair occludin tight junction of MTE monolayers. Laser scanning confocol imaging for occludin tight junction of MTE monolayers treated by 37°C **(a-c)**, 40°C for 120 min **(d-f)**, 500 μM acrolein (**g-i**), and the combination of 40°C and 500 μM acrolein **(j-l)**. Nuclei were labeled with hoechst (blue), and tight junctions were labeled with occludin (green). Scale bar: 10 μm. **(m-o)** Immunolocalization of claudin tight junction of normal MTE monolayer. Nuclei were labeled with hoechst (blue), and tight junctions were labeled with claudin (green). The expression of claudin was insufficient to be visualized. Scale bar: 50 μm. **(p)** Fluorescein isothiocyanate-dextran (4 kDa) permeability assay of MTE monolayers. MTE monolayers were treated by 37°C, 40°C for 120 min, 500 μM acrolein, and the combination of 40°C and 500 μM acrolein. Thermal stress and acrolein increased the permeability of fluorescein isothiocyanate-dextran. Student’s t-test. *\*\*\*P* < 0.001. *n* = 14.

**Supplemental data table 1.** Parameters when R_TE_ raw data were fitted by the ExpDecl function

| Group | P1 |  |  |  |  |  |  |  | P2 |  |  |  |
| --- | --- | --- | --- | --- | --- | --- | --- | --- | --- | --- | --- | --- |
| Sample number | 1 | 2 | 3 | 4 | 5 | 6 | 7 | 8 | 1 | 2 | 3 | 4 |
| $y_0$ | 0.34 | 0.46 | 0.52 | 0.097 | 0.30 | 0.35 | 0.36 | 0.42 | -0.89 | -1.08 | -0.97 | -0.55 |
| $A$ | 0.92 | 0.97 | 0.96 | 0.87 | 0.87 | 0.83 | 0.91 | 0.64 | 3.13 | 3.88 | 3.62 | 2.26 |

